# Molecular Understanding of Calorimetric Protein Unfolding Experiments

**DOI:** 10.1101/2021.08.10.455817

**Authors:** Joachim Seelig, Anna Seelig

## Abstract

Protein unfolding is a dynamic cooperative equilibrium between short lived protein conformations. The Zimm-Bragg theory is an ideal algorithm to handle cooperative processes. Here, we extend the analytical capabilities of the Zimm-Bragg theory in two directions. First, we combine the Zimm-Bragg partition function Z(T) with statistical-mechanical thermodynamics, explaining the thermodynamic system properties enthalpy, entropy and free energy with molecular parameters only. Second, the molecular enthalpy h_0_ to unfold a single amino acid residue is made temperature-dependent. The addition of a heat capacity term c_v_ allows predicting not only heat denaturation, but also cold denaturation. Moreover, it predicts the heat capacity increase 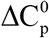 in protein unfolding. The theory is successfully applied to differential scanning calorimetry experiments of proteins of different size and structure, that is, gpW62 (62aa), ubiquitin (74aa), lysozyme (129aa), metmyoglobin (153aa) and mAb monoclonal antibody (1290aa). Particular attention was given to the free energy, which can easily be obtained from the heat capacity C_p_(T). The DSC experiments reveal a zero free energy for the native protein with an immediate decrease to negative free energies upon cold and heat denaturation. This trapezoidal shape is precisely reproduced by the Zimm-Bragg theory, whereas the so far applied non-cooperative 2-state model predicts a parabolic shape with a positive free energy maximum of the native protein. We demonstrate that the molecular parameters of the Zimm-Bragg theory have a well-defined physical meaning. In addition to predicting protein stability, independent of protein size, they yield estimates of unfolding kinetics and can be connected to molecular dynamics calculations.

## INTRODUCTION

Protein stability is an important issue in the development of pharmaceutical biologics. Thermodynamic aspects of protein folding have acquired significant practical importance because they provide the theoretical framework for rational protein design and protein modifications.[1] On the experimental level, differential scanning calorimetry (DSC) is the method of choice for thermodynamic studies of protein folding/unfolding equilibria.[2, 3] Analysis of DSC experiments with simple thermodynamic models has been key for developing our understanding of protein stability.[4] So far, the reversible denaturation reaction has been analyzed with a non-cooperative 2-state model.[5] However, its application is limited to small proteins with about 50 to 150 amino acids. Moreover, the free energy predicted by the 2-state model disagrees with the experimental finding obtained with differential scanning calorimetry.

The protein folding/unfolding transition is a dynamic equilibrium with many short-lived intermediates.[6] A multistate cooperative algorithm is therefore a physically more realistic alternative. We have shown with a variety of proteins that the cooperative multistate Zimm-Bragg theory is such a potential alternative. The theory yields a quantitated measure of cooperativity, is not limited in protein size and provides excellent simulations of protein unfolding thermodynamics.[7–14] Here we propose a significantly improved model where the Zimm-Bragg theory is combined with statistical-mechanical thermodynamics. First, we show that the macroscopic system parameters of unfolding, that is, enthalpy, entropy and free energy, can be traced back to simple molecular parameters of well-defined physical meaning. Second, the enthalpy h_0_ needed to unfold a single amino acid residue is supplemented with a heat capacity c_v_. A second unfolding transition at low temperature (cold denaturation) occurs and the heat capacity increase 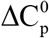 upon protein unfolding is also explained. The new model is validated by analyzing DSC measurements of different proteins, that is, lysozyme[10], the classic example of protein unfolding, gpW62[15], an ultrafast folding protein, mAb, a large monoclonal antibody[13], metmyoglobin[16], a protein exhibiting cold denaturation, and ubiquitin, an α-helical protein.[17] In the long-standing history of protein DSC measurements the heat capacity function C_p_(T) was mainly used to evaluate the enthalpy, whereas entropy and free energy were ignored, although both functions are easily derived from C_p_(T). Here we systematically evaluate entropy and free energy from C_p_(T) and analyze them with the multistate Zimm-Bragg theory. We show that the DSC observed temperature dependence of the free energy displays a trapezoidal shape with a zero free energy of the native state. This unique thermodynamic signature is precisely predicted by the Zimm-Bragg theory, but disagrees with the parabolic shape predicted by the 2-state model with a physically unrealistic positive free energy of the native state. Based on the molecular fit parameters at 20-90 °C the entropies of structurally different proteins were calculated at a denaturation temperature of 225 °C and were identical with the predictions of the Dynameomics Entropy Dictionary.[18]

## THEORY

### Zimm-Bragg theory extended to cold denaturation

Protein unfolding is a dynamic process in which individual amino acid residues flip from their native (n) to their unfolded (u) state. Rapid equilibria between many short-lived intermediates can be expected. An early example of cooperative unfolding is the α-helix-to-random coil transition of synthetic peptides described with the Zimm-Bragg theory.[19–21] The cooperative folding theory distinguishes between a growth process with an equilibrium constant q(T) and a nucleation step with an equilibrium constant σq(T) where σ is the cooperativity or nucleation parameter. Growth is defined as the addition of a new helical segment to an already existing α-helix. Nucleation is the formation of a new helical segment within an unstructured region. The steepness of the transition is determined by the cooperativity parameter σ, which is typically in the range of σ ~ 10^−3^ to 10^−7^. The cooperativity parameter of a non-cooperative system is σ = 1.

The central element of the Zimm-Bragg theory is the partition function Z(T) that collects all the energetic states of the folding process

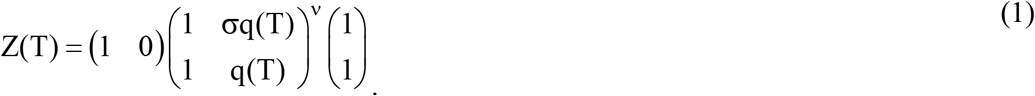

ν denotes the number of amino acid residues involved in unfolding. The equilibrium constant q(T) is given by

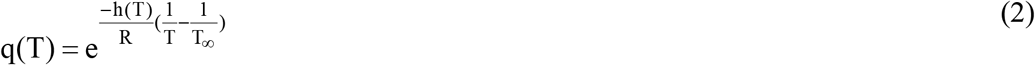

h(T) is the enthalpy needed to induce the n → u conformational change. Up until now, this parameter was assumed to be temperature-independent with h_0_ ~ 0.9 - 1.3 kcal/mol.[20, 22–25] Here we introduce a temperature-dependent unfolding enthalpy

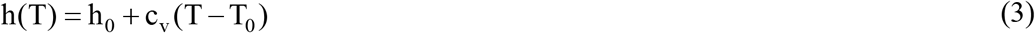

T_0_ is the midpoint temperature of heat-induced unfolding. The heat capacity c_v_ refers to the unfolding of a single amino acid residue. The reference temperature T_∞_ in equation (2) determines the position of the heat capacity maximum on the temperature axis. It is identical with T_0_ if the number of amino acid residues ν is much larger than σ^−1/2^. The steepness of the unfolding transition is determined by both σ and ν. The smaller the cooperativity parameter σ, the steeper is the unfolding transition. Conversely, the shorter the chain length ν, the broader is the unfolding transition. Short chains have a broad transition.[20]

The fraction of unfolded protein Θ_U_(T) is

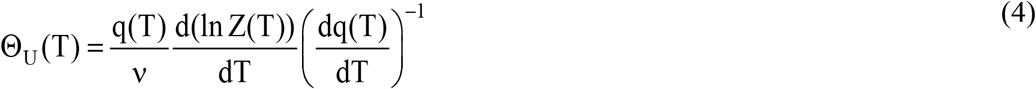

For a non-cooperative system with σ = 1 the multistate Zimm-Bragg theory reduces to a 2-state model.[14]

### Cold denaturation

The heat capacity term c_v_ has two consequences. First, c_v_ produces the heat capacity increase 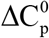 of the unfolded protein. Second, c_v_ leads to an additional transition at low temperature (cold denaturation). The exponent in the equilibrium constant q(T) (eq. (2)) has zeros at T_0_ and at h(T) = 0. The latter relation leads to

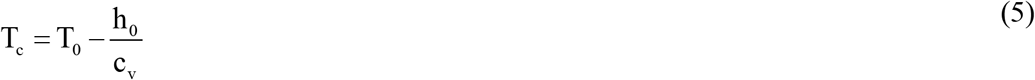

T_0_ is the midpoint of heat denaturation, T_c_ that of cold denaturation. The temperature difference between heat and cold denaturation is ΔT = h_0_/c_v_.

### Partition function and statistical-mechanical thermodynamics

We show that the thermodynamics of protein unfolding and, consequently, the DSC experiments can be simulated without macroscopic fit parameters. According to statistical mechanical thermodynamics the partition function Z(T) is the sum of all conformational energies and is sufficient to determine the thermodynamic system parameters, that is, the inner energy E(T), the entropy S(T), and the Helmholtz free energy F(T).[26, 27] The relevant equations are as follows

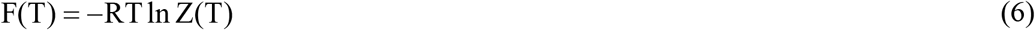

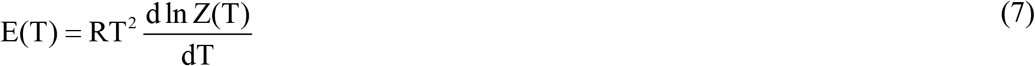

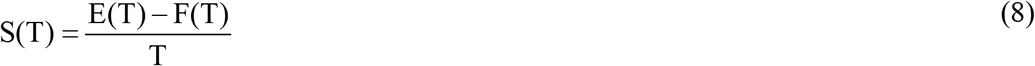

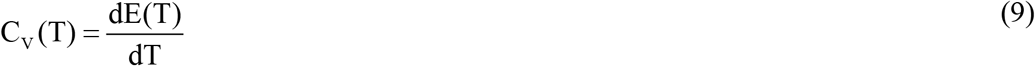

### Differential scanning calorimetry (DSC)

Differential scanning calorimetry (DSC) is the method of choice to determine the thermodynamic properties of protein unfolding. DSC measures the temperature course of the heat capacity C_p_(T) that includes not only the conformational enthalpy proper, but also the increase in heat capacity 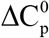 between native and denatured protein.[28, 29] Stepwise integration of C_p_(T) provides unfolding enthalpy, entropy, and Gibbs free energy:

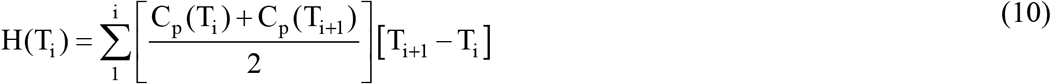

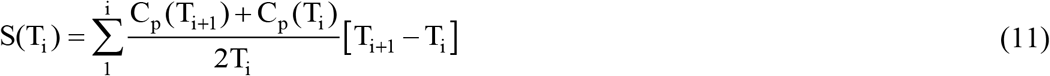

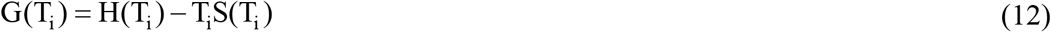

“It is clear that in considering the energetic characteristics of protein unfolding one has to take into account all energy which is accumulated upon heating and not only the very substantial heat effect associated with gross conformational transitions, that is, all the excess heat effects must be integrated”.[30] The DSC-measured thermodynamic parameters characterizing the total unfolding transition are denoted ΔH_cal_, ΔS_cal_ and ΔG_cal_.

DSC measurements are made at constant pressure. The volume changes in protein unfolding are very small and the following relations are valid without loss of accuracy. Heat capacity C_p_(T) ≅ C_v_(T), enthalpy H(T) ≅ inner energy E(T), entropy S_p_(T) ≅ S_v_(T), Gibbs free energy G(T) ≅ Helmholtz free energy F(T).

## Results

Our earlier simulations of DSC experiments with the Zimm-Bragg theory required molecular as well as macroscopic fit parameters. The extent of unfolding Θ_U_(T) was calculated with molecular parameters (h_0_, σ) and was then multiplied with macroscopic parameters, that is, the unfolding enthalpy ΔH_0_ and the heat capacity increase 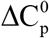. Here we eliminate ΔH_0_ and 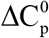.by combining the partition function Z(T) with statistical-mechanical thermodynamics (eqs. (6) – (9)). All macroscopic thermodynamic properties are now predicted exclusively with molecular parameters (h_0_, c_v_, σ, ν). Of particular interest is the free energy of unfolding G(T) ≅ F(T). Proposed as a parabolic function in the theory of the 2-state model, the DSC-accessible free energy G_cal_(T) (eq. (12)) appears to be completely ignored in applications of the 2-state model. In the following we compare proteins of different size and structure, which were carefully studied with DSC. The analysis of the C_p_(T) thermograms provides the caloric, model-independent results for enthalpy H_cal_(T), entropy S_cal_(T), and free energy G_cal_(T). The measured thermodynamic properties are then compared to the predictions by the cooperative Zimm-Bragg theory and the 2-state model.

### Lysozyme unfolding. DSC and molecular multistate partition function

Lysozyme is a 129-residue protein composed of ~25% α-helix, ~40% β-structure and ~35% random coil in solution at room temperature.[10] Upon unfolding, the α-helix is almost completely lost and the random coil content increases to ~60%. Thermal unfolding is completely reversible and lysozyme is the classical example to demonstrate 2-state unfolding.[2, 31, 32] Figure 1A shows the C_p_(T) thermograms of lysozyme unfolding with a resolution of 0.17°C [10] Unfolding takes place in the temperature range of 45 °C ≤ T ≤ 73 °C (midpoint temperature T_0_ = 61.7 °C). The unfolding enthalpy is ΔH_cal_ = 147 kcal/mol (eq. (10)) and the entropy ΔS_cal_ = 0.437 kcal/molK (eq. (11)) The enthalpy/entropy ratio ΔH_cal_/ΔS_cal_ = 335.5 K= 62.5 °C agrees with the midpoint temperature T_0_. The molar heat capacity of unfolded lysozyme increases by 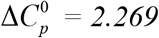 kcal/molK, in agreement with literature data.[28, 29, 32]

**Figure 1.**
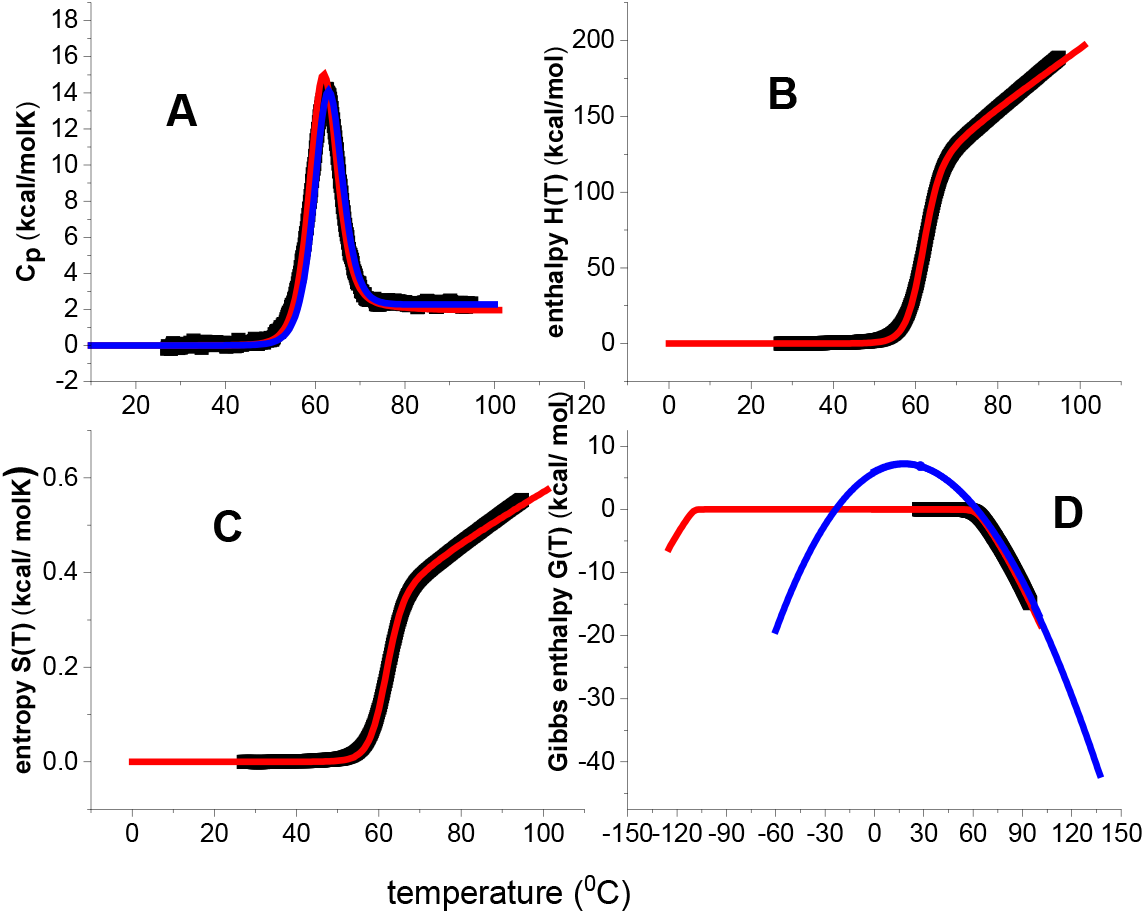
Differential scanning calorimetry of 50 μM lysozyme in 20% glycine buffer, pH 2.5. Black data points: DSC thermograms obtained with a heating rate of 1 °Cmin^−1^ and a step size of 0.17 °C. Red lines: simulations with the Zimm-Bragg theory with h_0_ =1.0 kcal/mol, c_ν_ = 6 cal/ molK, σ = 1.0×10^−6^, ν = 129. (A) Heat capacity C_p_(T). (B) Unfolding enthalpy H(T). (C) Unfolding entropy S(T). (D) Free energy of unfolding G(T). Blue line: ΔG(T) (eq.(13)) predicted by the 2-state model calculated with a conformational enthalpy ΔH_0_ = 107 kcal/mol and a heat capacity increase 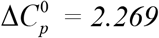 Kcal/mol. Data taken from reference.[10]

The high cooperativity of lysozyme unfolding (cooperativity parameter σ = 1.0×10^−6^) leads to a sharp unfolding transition, which is well approximated by a 2-state equilibrium. 2-state model and Zimm-Bragg theory both provide excellent simulations of the heat capacity C_p_(T) (fig. 1A).

Differences between the two models exist, however, in predicting the temperature dependence of the free energy. The 2-state model defines the free energy as

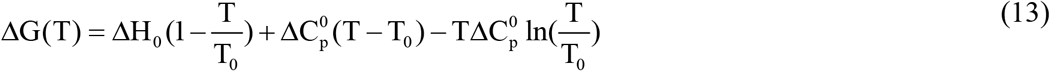

yielding the parabolic shape in figure 1D (blue line, same parameters as in Figure 1A). The free energy has a positive maximum of 7.18 kcal/mol for the native protein at 20 °C and becomes zero at T_0_ = 62°C, where lysozyme is 50% unfolded. However, this is not what is seen in the DSC experiment (fig. 1D, black data points). The free energy of the native protein is zero and becomes immediately negative upon unfolding. Between 27°C and T_0_ the DSC experiment reports a negative free energy change ΔG_cal_ = −0.7 kcal/mol, which increases rapidly beyond T_0_ to ΔG_cal_ = −6.06 kcal/mol at 73 °C (90% unfolding). The Zimm-Bragg theory (eq. (7)) reproduces precisely the DSC result (fig. 1D, red line). Extending this simulation to low temperatures yields cold denaturation at T_cold_ = −103 °C. The Zimm-Bragg theory thus predicts a trapezoidal shape of the free energy and the flat, near zero region extends over almost 140 °C. An experimental prove for the for the free energy trapeze is given in figure 4.

### gpW62, an ultrafast folding protein

gpW62 is a 62-residue α+β protein that belongs to a group of ultrafast folding proteins.[4, 15] gpW62 folding was investigated with a variety of techniques, including DSC. The heat capacity C_p_(T) (Figure 2A, data taken from reference[15]) is evaluated here according to eqs. (10) – (12). The midpoint temperature is T_0_ =67 °C. The unfolding enthalpy is ΔH_cal_ = 91.7 kcal/mol., measured between 37 °C and 102 °C. This is a large enthalpy for a 62-residue protein. The free energy change between native and unfolded gpW62 is ΔG_cal_ = −8.11 kcal/mol. The free energy per residue g_cal_ = −131 cal/mol is almost 3 times larger than that of lysozyme with g_cal_ = −47 cal/mol.

**Figure 2.**
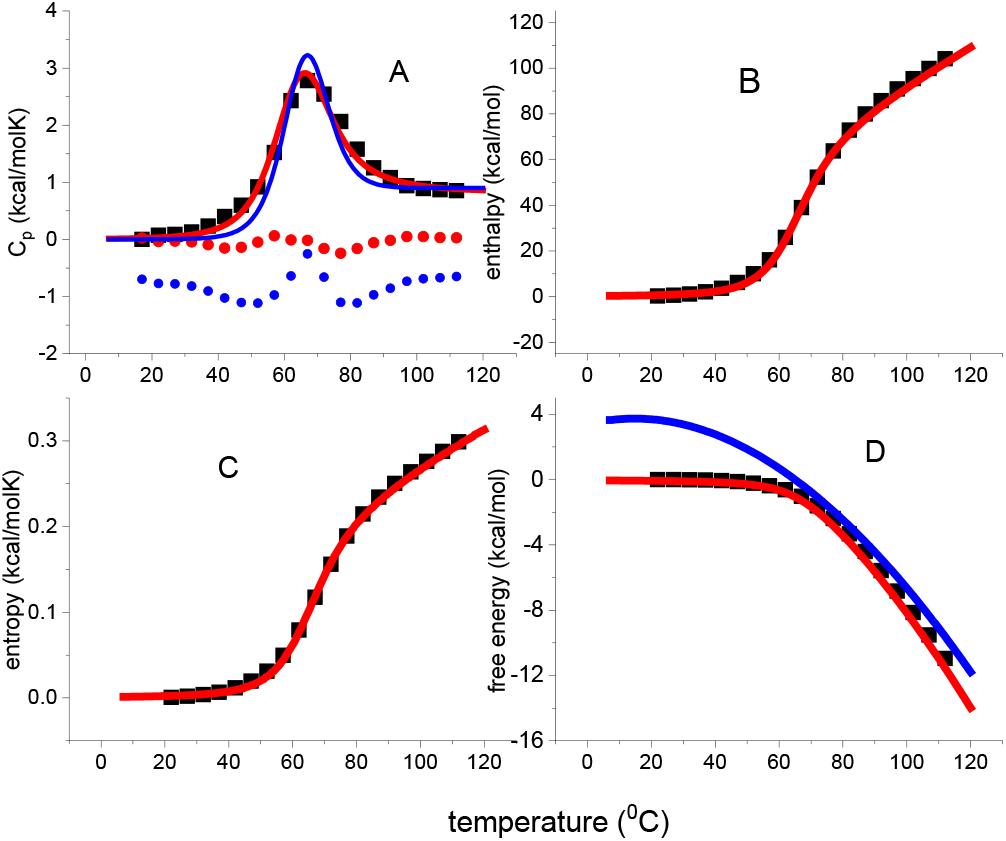
gpW62 DSC unfolding. (A) Heat capacity C_p_(T). Black data points taken from reference.[15] Data points in panels B-D calculated from data points in panel A with equations (10)–(12). Red solid lines: simulations with the Zimm-Bragg theory with h_0_ =1.26 kcal/mol, c_ν_ = 5 cal/molK, σ = 5.0×10^−4^, ν = 62 residues. Blue solid lines: 2-state model with ΔH_0_ = 49 kcal/mol and 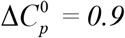 kcal/molK. The dotted lines in panel A are the differences between the experimental DSC data and the simulations. For better visibility the blue dotted line is shifted by −1 kcal/molK. (B) Unfolding enthalpy H(T). (C) Unfolding entropy S(T). (D) Free energy of unfolding G(T). Data taken from reference[15].

Several aspects of the gpW62 folding equilibrium are unusual. The cooperativity is low with σ = 5×10^−4^ and the unfolding extends over a broad temperature range of ΔT~65 °C. This may be compared to lysozyme with σ = 1×10^−6^ and ΔT~30 °C. As the transition is broad, the Zimm-Bragg theory provides a much better fit than the 2-state model (Figure 2A). The low cooperativity could facilitate fast folding by a low nucleation free energy (see Discussion). Fast folding of gpW62 could also be promoted by the large free energy change of the unfolding reaction. According to the thermodynamics of irreversible processes, the chemical reaction rate is proportional to the affinity, i.e. the free energy, of the reaction.[33–35] The ultra-fast folding of gpW62 could thus arise from the combination of a low nucleation energy and a large unfolding affinity.

### Monoclonal antibody mAb

The 2-state model works in a limited size range of about 100-200 kDa. No such size limitation exists for the Zimm-Bragg theory. This is demonstrated for the recombinant monoclonal IgG1 antibody mAb with a molecular weight of 143 kDa (~1280 amino acids).[13] The antibody is formed of two identical heavy chains of ~450 residues each and two identical light chains of ~200 residues. The heavy and light chains fold into domains of ~110 aa residues. The secondary structure of mAb is composed of 7-11% α-helix and 40-45% β-sheet.[36]

The DSC experiment (Figure 3A) reveals a pre-transition at 73 °C and a main transition at 85 °C. The unfolding enthalpy of the pre-transition is ΔH_cal_ = 290 kcal/mol involving ν_pre_ = ΔH_cal_/h_0_ = 263 amino acid residues. The main transition has an enthalpy of ΔH_cal_ ≃ 1000 kcal/mol with ν_main_ ≃880 amino acid residues. Taken together, pre- and main transition account for ~90% of all amino acid residues. A molecular interpretation of pre- and main transition based on the mAb structure has been given.[13] The pre-transition results from the unfolding of 2 C_H2_ domains, whereas the main transition represents the unfolding of 8 domains of the Fab fragment and 2 domains of the Fc fragment. In Figure 3 the pre-transition (green) and main transition (violet) are superimpose (red). The pre-transition is slightly less cooperative with σ = 5×10^−5^ than the main transition with σ = 2×10^−5^. The same molecular parameters h_0_ = 1.1 kcal/mol and c_v_ = 7.0 cal/molK were used in all simulations. The theory predicts the heat capacity increase upon unfolding as 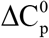 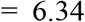 kcal/molK for the pre-transition and 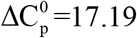 kcal/mol.K for the main transition, consistent with the number of amino acids involved. The DSC-measured temperature profile of the free energy follows the same pattern as observed for lysozyme and gpW62. The free energy of the native mAb is zero up to about 65 °C followed by a biphasic decrease. As shown in Figure 3D the contributions of the pre-transition (green) and the main transition (violet) are well separated. The multistate partition function Z(T) precisely predicts this biphasic behavior. The free energy per residue is g_cal_ = −30±1 cal/mol for both pre- and main transition and thus clearly smaller than those of lysozyme (−47 cal/mol) and gpW62 (−136 cal/mol). Neither the pre-transition nor the main transition can be simulated with the 2-state model.

**Figure 3.**
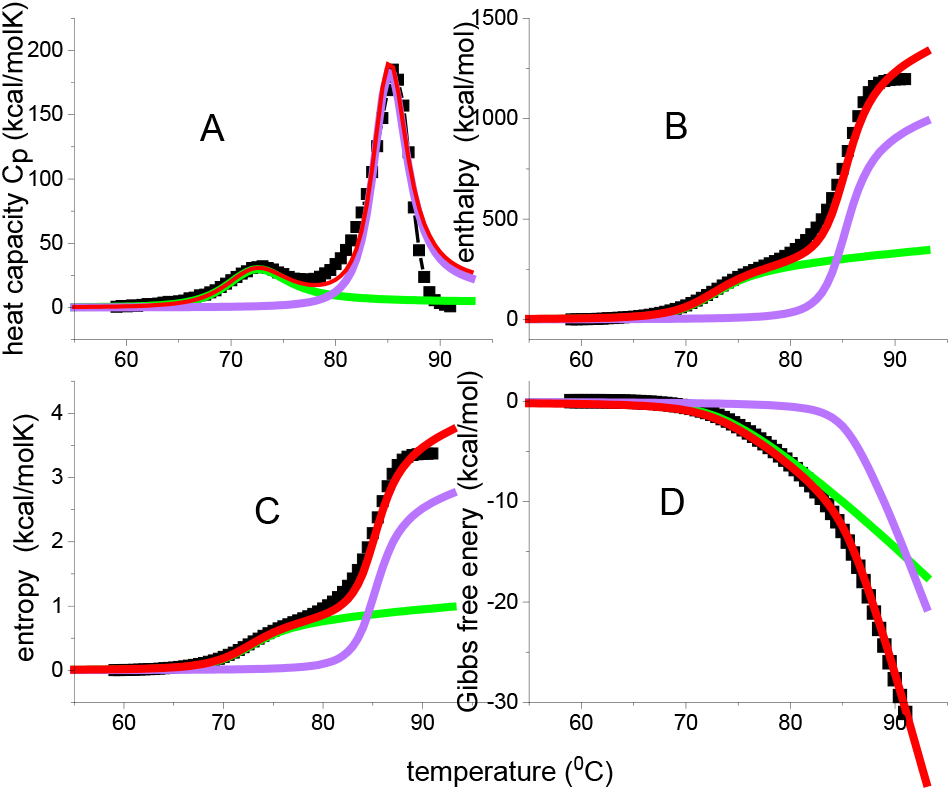
Thermal unfolding of monoclonal antibody mAb at pH 6.2. (■) DSC experiment. Solid lines are simulations with the Zimm-Bragg theory: green= pre-transition; violet= main transition; red= sum of pre- and main transition. (A) Molar heat capacity. (B) Unfolding enthalpy. (C) Unfolding entropy. (D) Free energy. h_0_ = 1.1 kcal/mol, c_ν_ = 7.0 cal/molK. Pre-transition: T_0_ = 73 °C, ν_pre_ = 263, σ = 5 × 10^−5^. Main transition: T_0_ = 85.4 °C, ν_main_ = 880, σ = 2 × 10^−5^. Data taken from reference.[13]

### Heat and cold denaturation of metmyoglobin

A protein that is stable at ambient temperature can be unfolded by heating or, less common, by cooling. The original Zimm-Bragg theory was modified here to include cold denaturation. Cold denaturation of most proteins occurs at subzero temperatures. Rather drastic conditions are needed to shift cold denaturation above 0 °C (e.g. addition of denaturants, low or high pH). DSC experiments reporting cold denaturation or at least partial cold denaturation are available for metmyoglobin[16], staphylococcus nuclease[37], β-lactoglobulin[38], streptomyces subtilisin inhibitor[39] or thioredoxin.[40]

Metmyoglobin (153 residues) consists of 8 α-helical regions connected by loops.[41] DSC at pH 4.1 (Figure 4A) displays heat denaturation at T_0_ = 69 °C and cold denaturation starting at 3 °C (data from Figure 13, reference [16]). Heat denaturation of the partially destabilised protein is characterized by ΔH_cal_ = 146 kcal/mol, ΔS_cal_ = 0.431 kcal/mol and ΔG_cal_ = −6.4 kcal/mol. The ratio ΔH_cal_/ΔS_cal_ = 339 K = 66 °C is consistent with the C_p_(T) maximum. Cold denaturation is predicted at Tcold=- −4.5 °C and at −3°C by the 2-state model. The Zimm-Bragg theory (fit parameters in table 1) provides a clearly better fit to C_p_(T) than the 2-state model (fit parameters ΔH_0_ = 112 kcal/mol, 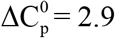 kcal/molK).

**Figure 4.**
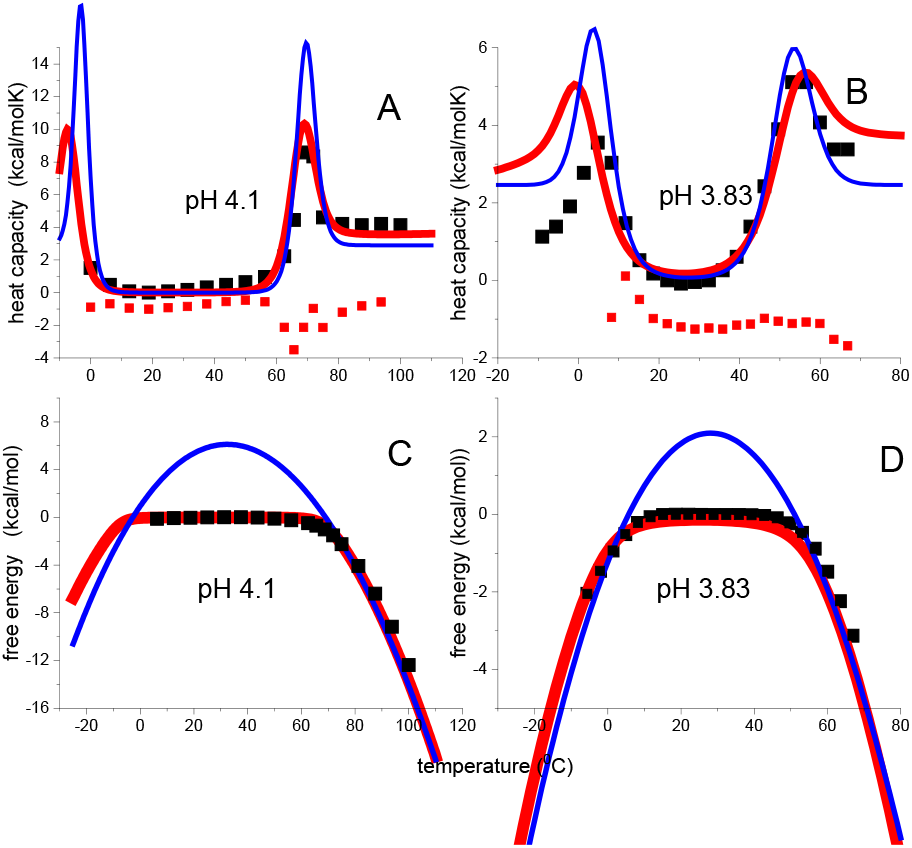
Unfolding of metmyoglobin at acid pH. (■) Experimental data taken from reerence[16]. Red lines: simulations with the cooperative Zimm-Bragg theory (fit parameters listed in table 1). 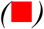 Differences between DSC data and the Zimm-Bragg simulation (shifted in 4B by –1 kcal/ molK for better visibility). Blue lines: 2-state model. (A) C_p_(T) at pH 4.1(Figure 13 in reference[16]). (B) C_p_(T) at pH 3.83 (Figure 12 in reference[16]). (C, D) Free energies calculated using the heat capacity data shown in panels 4A and 4B.

**Table 1.**
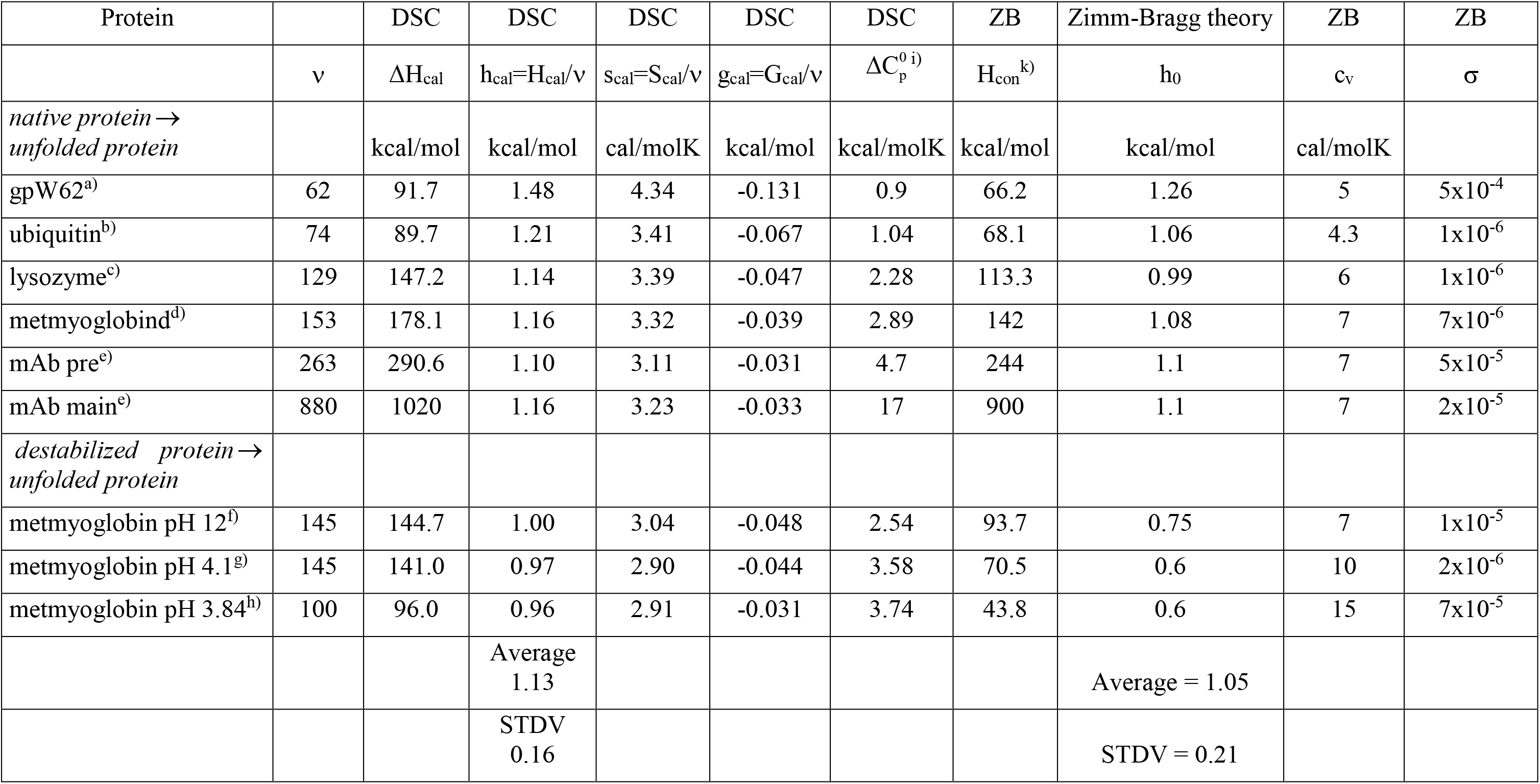
Thermal unfolding. Differential scanning calorimetry and parameters of the Zimm-Bragg theory. a) reference [15], fig. 4A (this work: figs. 2& 5 A); b) reference[17], Figure 1 (this work: Figure 5B) c)reference[10], Figure8 (this work: figs. 1 & 5C); d) reference[16], Figure3b, pH 10.7 (this work: Figure 5D); e) reference[13] (this work: Figure 3); f) reference[16], Figure 3b (no simulation shown); g) reference[16] Figure 13 (this work: figs. 4A&4C); h) reference[16] figure12 (this work: figs. 4B&4D); i). DSC-measured increase in molar heat capacity upon protein unfolding; k) conformational unfolding enthalpy proper, calculated with the Zimm-Bragg theory by setting c_v_ = 0.

At pH 3.83 metmyoglobin is even more destabilized and DSC reports two unfolding transitions with C_p_(T)-maxima at T_cold_ = 8 °C and T_0_ = 56.5 °C (Figure 4B). The enthalpy of heat unfolding is ΔH_cal_ = 96 kcal/mol and the entropy S_cal_ = 0.291. The ratio ΔH_cal_/ΔS_cal_ = 329.9 K = 56.9 °C agrees with T_0_. Cold denaturation is not the mirror image of heat denaturation as unfolding enthalpy and entropy are much smaller with ΔH_cold_ ~ 53.8 kcal/mol and ΔS_cold_ ~ 0.193 kcal/molK, respectively. The ratio ΔH_cold_/ΔS_cold_ = 274K = 1°C is different from T_cold_ = 8 °C. The low-temperature transition could be distorted in the DSC experiment. Again the Zimm-Bragg theory (fit parameters table 1) provides a better simulation than the 2-state model (fit parameters ΔH_0_ = 58 kcal/mol, 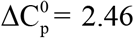 kcal/molK). . The free energy profile is displayed in Figure 4D. DSC yields a zero free energy for the native protein and negative free energies for heat and cold denaturation. This trapezoidal temperature profile is correctly predicted by the Zimm-Bragg theory. The specifics in the simulation of cold denaturation are a large heat capacity c_v_ and a small unfolding enthalpy h_0_ (cf. Table 1). In contrast, the 2-state model predicts a parabolic profile and assigns a large positive free energy to the native molecule.

Metmyoglobin at pH 3.83 was analyzed previously by a hierarchical algorithm defining a partition function in terms of multiple levels of interacting folding units.[42] The model reproduces an idealized, symmetrical shape of the heat capacity C_p_(T) and a parabolic free energy function.

Table 1 summarizes the DSC results and the Zimm-Bragg fit parameters of all proteins discussed. The table includes additional measurements of metmyoglobin at pH 10 and 12 and of ubiquitin[17] for which no simulations are shown.

## Discussion

Proteins in solution do not show a simple, 2-state equilibrium between a fully folded and a fully unfolded conformation. Depending on temperature, they form a complex mixture of many short-lived intermediates. Here we present a new model, which predicts the important thermodynamic functions enthalpy, entropy and free energy on the basis of molecular parameters only. The performance of the model is demonstrated by comparison with DSC experiments. The Zimm-Bragg theory provides excellent simulations of the temperature course of all thermodynamic functions reported by DSC. With this model we obtain insights into the cooperativity and dynamics of protein folding.

### Protein stability

The most relevant parameter of protein stability is the midpoint temperature of unfolding T_0_. Protein unfolding can be approximated by a first-order phase transition, and T_0_ is determined by T_0_ = ΔH_cal_/ΔS_cal_. ΔH_cal_ and ΔS_cal_ are the DSC-measured unfolding enthalpy and entropy, respectively. Minor changes in ΔH_cal_ or ΔS_cal_. produce distinctive shifts in T_0_. The ultrafast folder gpW62 (62 aa) and ubiquitin (74aa) are short proteins with almost identical unfolding enthalpies of 91.7 kcal/mol and 89.7 kcal/mol, respectively. Nevertheless, their transition temperatures are ~20 °C apart with gpW62 at 67 °C and ubiquitin at 89.5 °C. The difference is caused by the larger gpW62 entropy ΔS_cal_ = 0.269 kcal/molK compared to ΔS_cal_ = 0.25 kcal/molK of ubiquitin. The difference becomes even more obvious on a per residue basis with s_cal_=ΔS_cal_/ν= 4.34 kcal/mol for gpW62 and s_cal_= 3.41 kcal/mol for ubiquitin. Likewise, very small differences in enthalpy and entropy cause the 12 °C difference in the midpoint temperatures of the two mAb domains. *A priori* predictions of T_0_ therefore require highly precise MD calculations of unfolding enthalpy and entropy.

In a true first-order phase transition (e.g. melting of ice) the heat ΔH_cal_ is absorbed at the constant temperature T_0_ and the heat capacity is a sharp peak (singularity). In contrast, ΔH_cal_ in protein unfolding is absorbed over 20-60°C and the heat capacity C_p_(T) is a broad peak. In particular, and not generally recognized, the relation ΔH_cal_ = T_0_ΔS_cal_ is limited to the overall reaction, but not applicable to the measured heat H(T_0_) and entropy S(T_0_). Considering lysozyme as an example, the DSC-measured heat absorbed up to T_0_ is H(T_0_) = 63.4kcal/mol (eq. 10) and is less than half of the total heat of 147.2 kcal/mol). The corresponding entropy is S(T_0_) = 0.191 kcal/molK (eq. 11). As H(T_0_) < T_0_S(T_0_) this results in a negative free energy G(T_0_) = −0.677 kcal/mol (eq. 12), more realistic for a~50% unfolded protein than the zero free energy predicted by the 2-state model. Analogous results are obtained for all proteins discussed here.

The two hallmarks of the 2-state model, that is, the positive free energy of the native protein and the zero free energy of the 1:1 folded/unfolded mixture, are thus not confirmed by DSC. Instead, the free energy shows a trapezoidal shape that is precisely predicted by the new Zimm-Bragg folding model

### Molecular unfolding enthalpy h(T)

The Zimm-Bragg parameter h_0_ is an average value of all types of interactions, independent of specific conformations (α-helix, β-sheet, ionic interactions, etc.). h_0_ is close to the calorimetric average h_cal_ = ΔH_cal_/ν. Metmyoglobin is an α-helical protein and the average enthalpy of the native protein (pH 10) is h_cal_ = 178/153 = 1.16 kcal/mol while h_0_ = 1.08 kcal/mol (Figure 3 of ref[16], parameters listed in table 1, simulation not shown). For the ~50% α-helical apolipoprotein A1 the DSC result is h_cal_ = 1.08±0.07 kcal/mol and the Zimm-Bragg parameter h_0_ = 1.1 kcal/mol.[9] Lysozyme, a globular protein with mainly β-sheet structure, yields h_cal_ = 1.14kcal/mol and h_0_ = 0.99 kcal/mol (pH 2.5).[10] A comparison of a larger set of proteins has led to the conclusion that the Zimm-Bragg parameter is h_0_ = 1.1±0.2 kcal/mol.[10]

The enthalpy h_0_ is usually associated with the opening of an α-helix hydrogen bond.[20, 22–24] However, MD calculations have led to the conclusion that “hydrogen bond formation contributes little to helix stability […] The major driving force for helix is associated with interactions, including van der Waals interactions, in the close packed helix conformation and the hydrophobic effect”.[43] This is supported by experimental results obtained with short alanine-based peptides, where hydrophobic interactions play the dominant role in stabilizing isolated α-helices.[25]

### Cold denaturation

Cold denaturation has been proposed as a tool to measure protein stability.[44] The heat capacity c_v_ is a new parameter in the Zimm-Bragg theory leading to a second unfolding transition at low temperature. The temperature difference between heat and cold denaturation is given by ΔT= h_0_/c_v_ (eq. (5)). Cold denaturation near ambient temperature thus requires a small h_0_ and a large c_v_.. This is confirmed by metmyoglobin at pH 4.1 and 3.83, where h_0_ is distinctly reduced to h_0_ = ~0.6 kcal/mol where c_v_ = 10-15 cal/molK, that is, twice as large as that of the native protein (cf. table 1). Proteins with h_0_/c_v_ ≥ 100 K display cold denaturation at very low sub-zero temperatures.

### Molecular unfolding entropy

Molecular dynamics (MD) calculations consider all possible degrees of dihedral freedom of each amino acid residue in sampling the conformational space. In contrast, the Zimm-Bragg theory is a algorithm that differentiates only between folded and non-folded amino acid residues, independent of the specific conformation. However, the theory makes predictions, solidly based on experimental data, which can be compared to MD calculations. The temperature course of the entropy is a good example. The entropy S(T) can be calculated with the Zimm-Bragg theory according to eq.(8). The excellent agreement with the experimental data is displayed in Figures 1–3. In Figure 5 we have repeated these calculations, including also ubiquitin, for the much larger temperature range of 298K-498K as this is the temperature interval of the “Dynameomics Entropy Dictionary”.[18]

**Figure 5.**
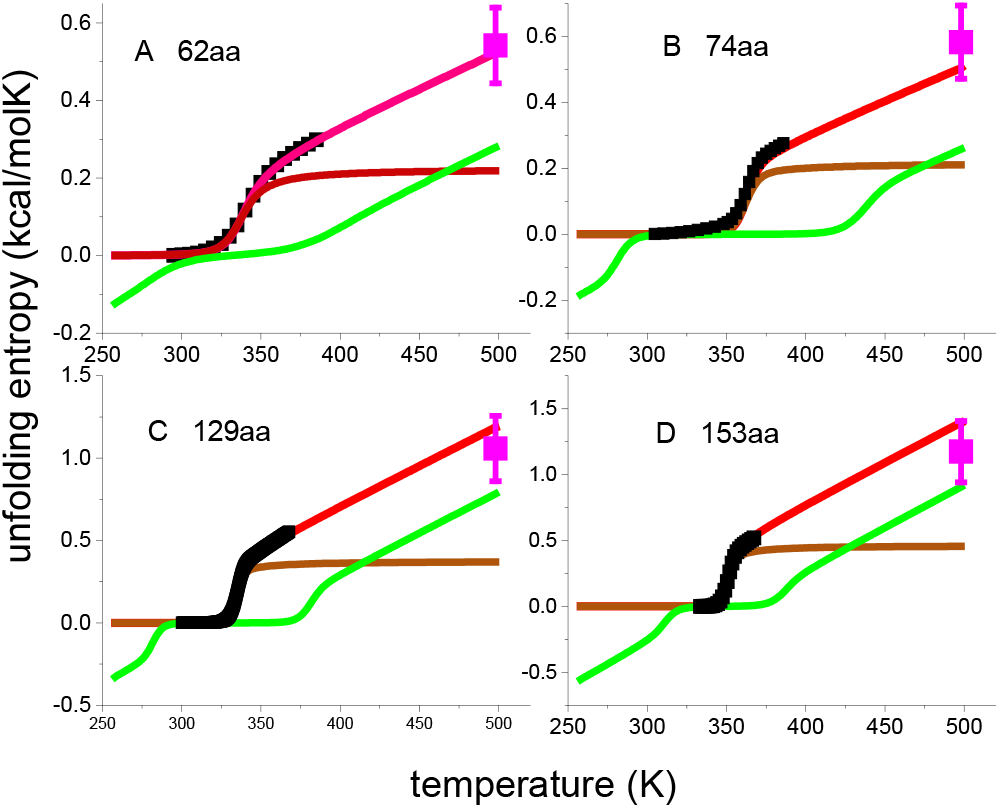
Unfolding entropy. (■)Experimental data.. 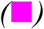 Dynameonics Entropy Dictionary[18]. (A) gpW62.[15].(B) Ubiquitin.[17] (Figure 1 in.[17]) (C) Lysozyme[10] (D) Metmyoglobin.[16](Figure 3 in ref. [16], pH 10). Red lines: Zimm-Bragg total entropy. Brown lines: conformational entropy proper. Green lines: contribution of the heat capacity term cv, Zimm-Bragg parameters listed in table 1.

By averaging some 800 MD calculations the Dynameomics Entropy Dictionary provides the unfolding entropies of all amino acids when heated from the native state (298K) to the fully denatured state (498K). Using table 2 in reference[18], we calculated the MD unfolding entropies at 498K for the specific amino acid sequences of the 4 proteins in Figure 5. The results are shown in Figure 5 by the magenta data points at 498K with the error bars also taken from reference.[18] In parallel, the Zimm-Bragg simulations were extended to 498K with the same parameters as deduced from C_p_(T) at low temperature. This extrapolation is in excellent agreement with the Dynameomics Entropy Dictionary and supports the relevance of the molecular parameters of the Zimm-Bragg theory in protein unfolding.

The entropy S(T)(red) can be divided into the conformational entropy proper (brown line, h_0_ = const., c_v_ = 0) and the contribution of the heat capacity term (green, h_0_ = 0, c_v_=const.). h_0_ determines S(T) up to the end of the conformational transition where Θ_U_ ~ 0.85-0.9. However, it takes another temperature increase of more than 100 °C to reach complete denaturation.

The average entropy is s_cal_ = ΔS_cal_/ν = 3.25±0.25 cal/molK per residue (table 1) and involves at least 3 single bonds. The entropy per single bond is ~1.1 cal/molK and can be compared with other phase transitions. The solid-fluid transition of long chain paraffins yields a much larger entropy of 1.8-2.7 cal/molK per C-C bond. The gel-to liquid crystal transition of phospholipid bilayers leads 1.25 - 1.6 cal/molK per C-C bond (table 2.7, p. 47 in reference[45]). As judged by the small entropy of 1.1 cal/molK the unfolded proteins are still characterized by a restricted motion of their molecular constituents. For metmyoglobin it was noted that “the unfolded state retains some residual ellipticity, which may be caused by the fluctuating α-helical conformation of the unfolded polypeptide chain”.[16] Likewise, the combination of FRET, NMR and SAX techniques has revealed residual structure in denatured ubiquitin.[46]

### Protein cooperativity

The folding/unfolding transition of proteins is a cooperative process. The Zimm-Bragg theory provides a quantitated measure of cooperativity. In fact, the cooperativity parameter σ plays an important role in the energy and kinetics of the folding process as it determines the free energy to start a new fold within an unfolded region (nucleation).[47] The corresponding free energy is

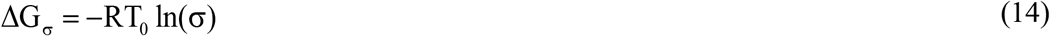

A large σ corresponds to low cooperativity and to a small nucleation energy ΔG_σ_. gpW62 has a cooperativity parameter σ = 5×10^−4^ and hence a nucleation energy of ΔG_σ_ = 5.13 kcal/mol. In contrast, lysozyme unfolding is highly cooperative with σ = 1×10^−6^ and ΔG_σ_ = 9.21 kcal/mol. The low gpW62 nucleation energy makes gpW62 folding easier than that of lysozyme. If ΔG_σ_ is assumed to be correlated with the kinetic activation energy, then gpW62 folding should be ~500 times faster than that of lysozyme. In infra-red T-jump experiments of gpW62 the return to equilibrium followed a single exponential with a relaxation time of τ = 15.7 μs at 57 °C.[15] In contrast, lysozyme was found to fold in a two-step mechanism with a slow nucleation (τ = 14 ms) followed by a fast growth step (τ = 300 μs) at room temperature.[48]

### Free energy of unfolding

The free energy of unfolding ΔG_cal_ scales with the size of the protein and a per residue free energy g_cal_ = ΔG_cal_/ν is better suited for comparative purposes. According to the Zimm-Bragg theory g_cal_ can be approximated by

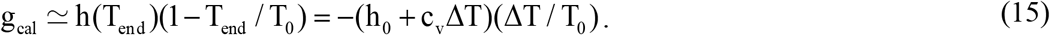

T_end_ denotes the temperature at the end of the conformational transition and ΔT = T_end_−T_0_ is the half-width of the transition. The approximation eq. (15) agrees within 2% with the DSC measurement for the proteins listed in table 1. The free energy per amino acid residue varies between g_cal_ = −131 cal/mol for gpW62 to g_cal_ = −33 cal/mol for mAb. In parallel, the width of the transition decreases from 65 °C for gpW62 to 20 °C for mAb (large domain). An approximately linear relationship between gcal and ΔT is predicted by eq. (15) and is confirmed by DSC, that is, a large g_cal_ correlates with a broad transition. Considering the three parameters T_0_, g_cal_, and ΔT, gpW62 is the least stable protein discussed here.

## Conclusion

The protein folding/unfolding transition is a highly cooperative process which cannot be adequately simulated by the non-cooperative 2-state model. A multistate cooperative model is provided by the Zimm-Bragg theory. Here we combined the partition function of the Zimm-Bragg theory with statistical-mechanical thermodynamics. The model predicts the DSC-measured enthalpy, entropy, and free energy of protein unfolding with molecular parameters, which have well-defined physical meanings. We analyzed the DSC thermograms of proteins of different chain length and structure. We show that the temperature profile of the free energy is characterized by a trapezoidal shape. The new model is in excellent agreement with this experimental finding. In contrast to the 2-state model that postulates a parabolic shape. The present model reveals whether a protein is a fast or a slow folder and predicts heat as well as cold denaturation. The results of the new model can be compared to molecular dynamics calculations. Using the molecular parameters derived from DSC the entropy at complete unfolding at 498K was calculated for four different proteins. The results are in excellent agreement with the predictions of the Dynameomics Entropy Dictionary. A detailed theoretical discussion of the free energy in thermal and chemical protein unfolding is available.[49]

